# Short-term Ketogenic Diet Induces a Molecular Response that is Distinct from Dietary Protein Restriction

**DOI:** 10.1101/2021.12.19.473355

**Authors:** Krystle C. Kalafut, Sarah J. Mitchell, Michael R. MacArthur, James R. Mitchell

**Affiliations:** Harvard T.H. Chan School of Public Health, Department of Molecular Metabolism, Boston, MA, USA; Department of Health Sciences and Technology, Swiss Federal Institute of Technology (ETH) Zurich, Zurich, Switzerland

**Keywords:** ketogenic, dietary restriction, protein restriction, carbohydrate, protein, FGF21

## Abstract

There is increasing interest in utilizing short-term dietary interventions in the contexts of cancer, surgical stress and metabolic disease. These short-term diets may be more feasible than extended interventions and may be designed to complement existing therapies. In particular, the high-fat, low-carbohydrate ketogenic diet (KD), traditionally used to treat epilepsy, has gained popularity as a potential strategy for weight loss and improved metabolic health. In mice, long-term KD improves insulin sensitivity and extends lifespan and healthspan. Dietary protein restriction (PR) causes increased energy expenditure, weight loss and improved glucose homeostasis. Since KD is inherently a low-protein diet (10% of calories from protein vs. 20% in control diet), here we evaluated the potential for mechanistic overlap between PR and KD via activation of a PR response. Mice were fed control, protein-free, or one of four ketogenic diets with varying protein content for 8 days. PF and KD diets both decreased body weight, fat mass, and liver weights, and reduced fasting glucose and insulin levels, compared to mice fed the control diet. However, PF and KD differed with respect to insulin tolerance and hepatic insulin sensitivity, which were increased in PF-fed mice and impaired in KD-fed mice relative to controls. Furthermore, contrary to the PF-fed mice, mice fed ketogenic diets containing at least 5% protein did not increase hepatic *Fgf21* or brown adipose *Ucp1* expression. Interestingly, mice fed KD lacking protein demonstrated greater elevations in hepatic *Fgf21* than mice fed a low-fat PF diet. To further elucidate potential mechanistic differences between PF and KD diets and the interplay between dietary protein and carbohydrate restriction, we conducted RNA-seq analysis on livers from mice fed each of the six diets and identified distinct gene sets which respond to dietary protein content, dietary fat content, and ketogenesis. We conclude that KD with 10% of energy from protein does not induce a protein restriction response, and that the overlapping metabolic benefits of KD and PF diets occur via distinct underlying mechanisms.

## 1 Introduction

The ability of dietary interventions to improve metabolic health and stress resistance has been extensively demonstrated in animal models since the first studies on caloric restriction in the early 1900’s (1–3). Many recent studies have focused on dietary restriction of specific macronutrients, which in some cases has been shown to have similar benefits as total caloric restriction. Both the high-fat, low-carbohydrate ketogenic diet (KD) and protein restricted (PR) diets improve metabolic health and stress resistance in mice (4–7). However, whether there is an overlapping mechanism that underlies the benefits of these diets remains to be determined.

Long-term feeding of both KD and PR diets improves metabolic health in mice. PR diets with a low protein-to-carbohydrate ratio improve measures of cardiometabolic health and glucose tolerance in mice and are associated with lower body weight, and reduced plasma lipids and insulin (8–10). Similarly, long-term KD reduces body weight gain and improves insulin and glucose tolerance in mice when started early in life (11). Although the metabolic phenotypes of extended KD and PR interventions are similar, studies evaluating short-term KD (≤5 weeks) found impaired hepatic insulin resistance, despite significantly reduced plasma glucose and insulin (12–14). This contrasts with PR diets which show profound improvements in glucose and lipid homeostasis after just 7 days (15).

Clinically, both KD and PR have a history of long-term use in the treatment of idiopathic epilepsy in adults and children (16–21) and renal failure (22–26), respectively. In addition to effectively treating these conditions, both dietary interventions were observed to improve cardiometabolic parameters in human patients. KD has been shown to cause significant weight loss and reduction in cardiovascular risk factors, including blood pressure, and serum triglycerides and insulin, in non-epileptic humans with obesity (27). PR has been shown to substantially improve glucose and insulin homeostasis in humans with and without diabetes (28–31). In addition, high protein intake is associated with increased overall mortality and mortality from cancer or diabetes in individuals under the age of 65 (32).

Recently, the therapeutic value of short-term or cyclic KD and PR are being investigated for a variety of indications aside from epilepsy and renal insufficiency. Studies in mice have demonstrated that 10 days of KD-feeding can improve the efficacy of PI3K inhibitors in xenograft models (33), and short-term KD was recently shown to impact xenograft tumor growth through alterations in metabolic signaling and nutrient availability (34). Previous pilot studies have demonstrated that KD is safe and feasible in patients with glioblastoma and other advanced cancers (35–37), and there is an ongoing pilot study to evaluate the tolerability of KD in endometrial cancer patients (NCT03285152). Short-term KD has also been proposed to protect against adipose tissue inflammation in mice (38). In addition, a recent report showed that one-week cycles of KD alternated with a control diet reduces midlife mortality and improves measures of healthspan and memory in mice (39). The fasting mimicking diet (FMD) combines calorie restriction and PR and is being tested as an adjunct to chemotherapy in multiple cancer types (40, 41). Short 4- or 5-day cycles of FMD have also been shown to improve measures of cardiometabolic health in humans (42), and to improve late-life health, included delayed cancer incidence, in mice (43). Preconditioning with short-term PR also effectively protects against ischemic surgical stress in mouse models (4, 44) and a combination of calorie restriction and PR has been shown to be safe in humans prior to surgery (45, 46). Given the recent clinical interest in shorter-term KD and PR diets, it is important to understand the mechanism by which these interventions improve health and stress resistance to fully harness their therapeutic potential.

Both KD and PR are known to induce numerous metabolic adaptations that may contribute to the observed benefits. PR is associated with increases in energy expenditure and thermogenesis, as well as increased circulating FGF21, a hormone that regulates energy balance by acting on various tissues (47–50). KD is associated with improvements in lipid homeostasis, including upregulation of fatty acid oxidation and fat utilization and decreased lipid synthesis (14, 51). Interestingly, clinical KD diets used in humans are also restricted in protein content, with 5-10% of calories coming from protein compared to 15-20% in normal diets. This raises the question of whether overlapping molecular mechanisms related to reduced dietary protein content underlie the improvements in metabolic health and longevity in response to KD and PR.

The goal of this study was to compare the molecular signatures of mice fed KD to those of mice fed a protein-free (PF) diet. In addition to comparing low-protein/high carbohydrate (PF) to high fat/low carbohydrate ketogenic diet (KD), the protein content of KD was further titrated between 0-10% to evaluate the contribution of dietary protein to the KD phenotypes. After 1 week, PF and KD diets both decreased body weight, fat mass, and liver weights, and reduced fasting glucose and insulin levels, compared to mice fed the control diet. Contrary to the PF-fed mice, KD-fed mice did not increase markers of the physiological response to protein restriction, including hepatic *Fgf21* or BAT *Ucp1* expression. PF and KD also differed with respect to glucose and insulin tolerance and hepatic insulin signaling potential, which were all increased in PF-fed mice and impaired in KD-fed mice relative to controls. The intermediate diets revealed that increasing the protein content of the KD diet to 20% of energy impairs ketogenesis, as has been previously reported, but that replacing the protein in KD with carbohydrate does not. Liver transcriptomics further illuminated potential shared and independent mechanisms underlying the metabolic adaptations to carbohydrate and protein restriction, including shared alterations in fatty acid metabolism, but distinct changes in amino acid metabolism in PR and nucleotide metabolism in KD. We conclude that KD does not induce a robust protein restriction response, and that the overlapping metabolic benefits of KD and PF diets may be explained by distinct underlying mechanisms.

## 2 Methods

### 2.1 Mice

16-week-old male B6D2/F1 hybrid mice were purchased from Jackson Labs (strain no. 100006). Mice were acclimated to the Harvard T.H. Chan School of Public Health mouse facility for two weeks prior to experiments and were fed a standard chow diet during this acclimation period (Purina, 5053-PicoLab Rodent Diet 20). Mice were allowed ad libitum access to food and water unless otherwise noted. Mouse numbers were n=4 per diet group for the GTT, ITT, and insulin injection experiments, and a separate cohort of n=6 per diet group for all other analyses, with 2 mice co-housed per cage. Mice were maintained at 22°C with 12-hour light-dark cycles and 30-50% relative humidity. Mice were housed in a Specific Pathogen Free (SPF) facility, as determined by screening of sentinel animals at monthly intervals. Mice were fasted for six hours prior to sacrifice and tissue collection. All procedures were approved by the Harvard Institutional Animal Care and Use Committee.

### 2.2 Diets

Semi-purified diets were prepared using combinations of soluble protein-free base mixtures D12450Spx or D100070801Lpx (Research Diets, **Tables S1-2**), cocoa butter, casein, cystine and sucrose (**Fig. S1B**). Mixtures were combined with an equal mass of 2% agar in water to form a solid gel diet. Diets had macronutrient compositions and energy densities as listed in Supplemental Figure S1A. Food intake was measured daily at approximately ZT10.

### 2.3 Body Composition

Body mass was determined by daily measurement at approximately ZT10. Lean and fat mass were measured in awake mice using an EchoMRI analyzer system.

### 2.4 Serum Measurements

Mice were fasted for 6 hours prior to measurements and tail vein blood collection. Glucose was measured using a Clarity BG1000 handheld glucometer. Serum insulin was measured by ELISA following manufacturer protocol (Crystal Chem, #90080). Ketones were measured using a Nova Max Plus Glucose/Ketone handheld meter with Nova Max Ketone Strips. Serum FGF21 was measured by ELISA following manufacturer protocol (R&D Systems, #MF2100).

### 2.5 Glucose and Insulin Tolerance Tests

Metabolic assessments were performed in a separate cohort of mice fed either CTL, PF, or KD10P diets (n=4 per diet). An oral glucose tolerance test (OGTT) was performed on day 6 and an insulin tolerance test (ITT) was performed on day 8. For OGTT, mice were fasted for 6 hours and then 30% D-glucose (Sigma Aldrich) in sterile water was administered by oral gavage at 2g/kg. Blood glucose was measured from a small tail nick prior to (time 0) and at 15-, 30-, 60- and 120-minutes post-gavage using a handheld glucometer (Clarity BG1000). Blood was collected via tail vein at all timepoints, and serum was generated for insulin measurement.

For ITT, mice were fasted for 4 hours then injected intraperitoneally with 1.5IU/kg. Blood glucose was measured from a small tail nick prior to (time 0) and at 15-, 30-, 60- and 120-minutes post-injection as above.

### 2.6 Insulin Signaling

Mice fed CTL, PF, or KD10P diets for 10 days (n=4 per group) were fasted for 6 hours and then injected with 1.5 IU/kg insulin or vehicle for 20 minutes. Liver tissue was collected and flash frozen prior to analysis.

### 2.7 Immunoblotting

Flash-frozen liver tissue was homogenized and lysed in RIPA buffer (50mM Tris-Cl pH 7.4, 150mM NaCl, 1% IGEPAL, 0.5% sodium deoxycholic acid, 0.1% SDS, 1mM EDTA, 10mM NaF, 10mM sodium pyrophosphate, 1mM β-glycerophosphate, and 1mM sodium orthovanadate, Sigma protease inhibitor P8340). Protein quantification was performed using a BCA assay kit (Thermo Scientific) and equal amounts of protein were separated by SDS-PAGE, transferred to nitrocellulose membranes, and immunoblotted with indicated primary antibodies. Primary antibodies: p-IRS1 (CST, 2381), IRS (CST, 2382), p-FoxO1 (CST, 9464), FoxO1 (CST, 9462), p-Akt (CST, 4060), Akt (CST, 4691), p-rpS6 S235/36 (CST, 2211), rpS6 (CST, 2217), and Actin (CST, 4967). Secondary antibodies: anti-rabbit IgG, HRP-linked (CST, 7074), anti-mouse IgG, HRP-linked (CST, 7076). Immunoblots were developed by ECL (West Pico or Femto, Thermo Scientific).

### 2.8 RT-qPCR

RNA was isolated from flash-frozen liver or brown adipose tissue by homogenization in TRIzol Reagent (Thermo Fisher) followed by chloroform extraction and isopropanol precipitation. The concentration of RNA was determined using a Nanodrop Spectrophotometer. RNA (1μg) was reverse transcribed using the Advanced cDNA Synthesis Kit (Bio-Rad). qPCR was performed using Power Up SYBR green (Bio-Rad) with duplicate technical replicates using the QuantStudio 5 Real-Time PCR system. ΔΔCt values were normalized to actin and relative expression was plotted. Primer sequences were: Asns forward: GCAGTGTCTGAGTGCGATGAA, Asns reverse: TCTTATCGGCTGCATTCCAAAC, Fgf21 forward: CTGCTGGGGGTCTACCAAG, Fgf21 reverse: CTGCGCCTACCACTGTTCC, Ucp1 forward: AGGCTTCCAGTACCATTAGGT, Ucp1 reverse: CTGAGTGAGGCAAAGCTGATTT.

### 2.9 RNA-Seq

Livers were collected from euthanized mice and immediately flash frozen in liquid nitrogen and stored at −80°C until analysis. Livers were homogenized using a handheld homogenizer and RNA was extracted using a Qiagen RNeasy Plus Mini Kit (Qiagen #74134). The concentration and purity of RNA was determined using a Nanodrop Spectrophotometer and confirmed using an Agilent 2100 Bioanalyzer. Libraries were prepared using the Illumina TruSeq Stranded Total RNA Sample Preparation protocol. RNA was sequenced on an Illumina NovaSeq 6000 with 20 million paired end reads (150bp length) per sample.

Reads were aligned to the mouse GRCm38.p6 assembly using the align function and annotated using the featureCounts function from the Rsubread package (version 2.3.7). Differential expression analysis was performed using the edgeR (3.30.3) and limma (3.44.3) packages. Gene symbols were mapped to Entrez IDs using the mapIds function from the AnnotationDbi package (1.51.1). Cluster analysis was performed using the degPatterns function from the DEGreport (1.24.1) package. Normalization was performed using the trimmed mean of M-values method as implemented in the calcNormFactors from edgeR. Data were modeled and differential expression was determined using the limma voom pipeline to generate linear models with empirical Bayes moderation. Differential expression was determined using a Benjamini-Hochberg adjusted p value less than 0.05. Once differentially expressed genes or gene clusters were determined, gene set enrichment analysis was determined using the enrichKEGG function from the clusterProfiler package (3.16.0) (52). Weighted gene correlation network analysis was performed using the WGCNA package (1.70-3) (53). Transcription factor (TF) binding analyses were performed using CiiiDER (54).

### 2.10 Statistics

Statistical analyses were performed in Prism (version 8, GraphPad Software) and mean values were plotted with error bars representing standard deviations. One-way ANOVAs were followed by either Tukey’s post-hoc test or Dunnett’s post-hoc test. Transcriptomic data were analyzed using R (version 4.0.2) and multiple comparisons in transcriptomic data were corrected using the Benjamini-Hochberg false detection rate correction.

## 3 Results

### 3.1 Short-term dietary protein or carbohydrate restriction reduces body weight and adiposity in mice

In order to evaluate the contribution of protein restriction to the metabolic and molecular adaptations elicited by KD, mice were fed six distinct diets: Low-fat control containing 20% of energy from protein (CTL) or 0% of energy from protein (PF), classical KD containing no carbohydrate and 10% energy from protein (KD10P), modified KD for which half or all protein content was replaced with carbohydrate (KD5P or KD0P), or KD with protein content normalized to the control diet (KD20P) (**Fig. 1A; Fig. S1, A-B**).

**Figure 1.**
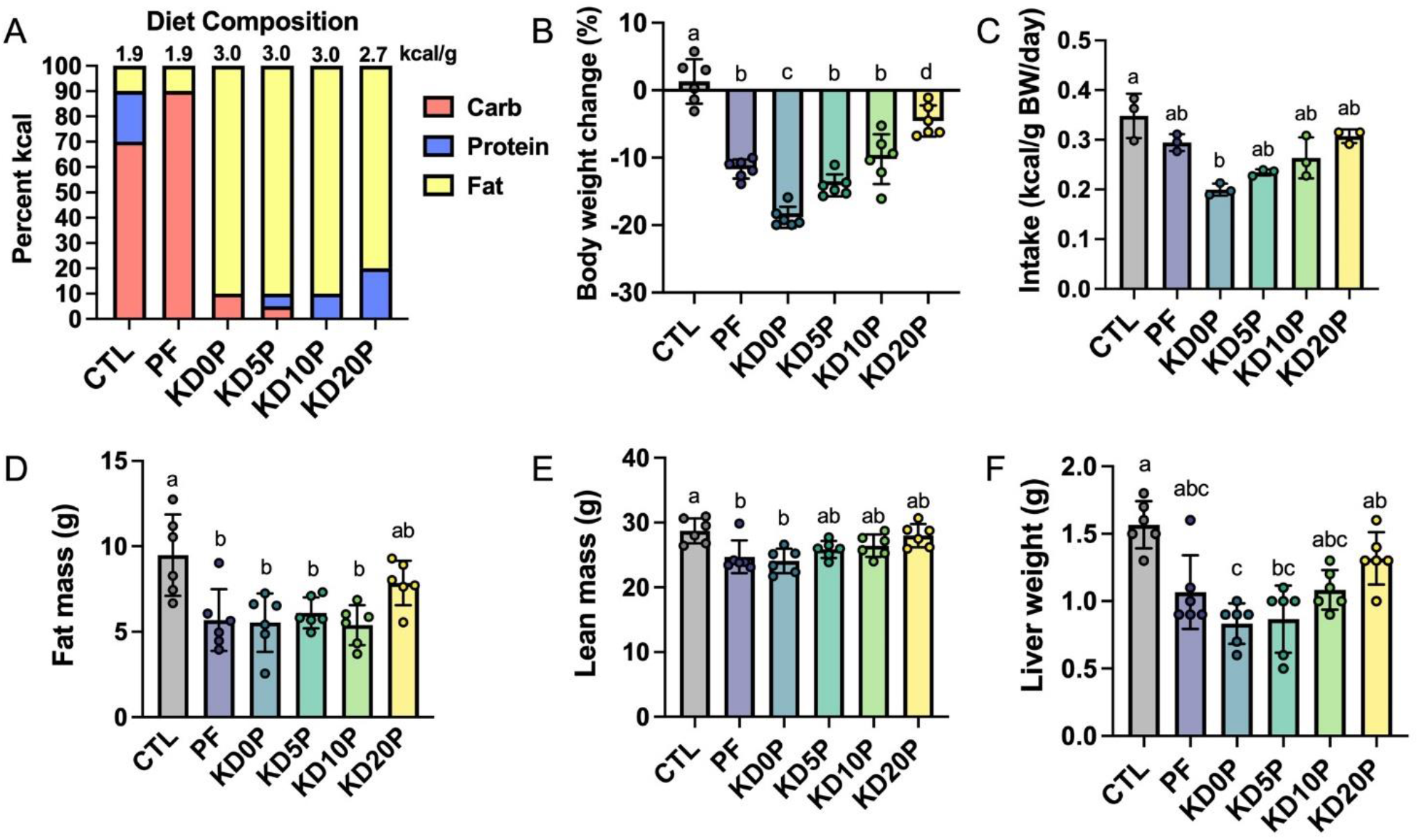
Short-term dietary protein or carbohydrate restriction alters body composition. Male B6D2F1 mice were fed one of six diets for 8 days (n=6 per diet) with varying macronutrient compositions as depicted in **(A)**. The energy density of each diet is listed above the associated bar. **(B)** Changes in body weight after 8 days of diet shown as the percent of the initial weight. **(C)** Average energy intake across 8 days of diet feeding. Intake was normalized to the summed body weight of 2 mice co-housed per cage and presented per cage. Fat mass **(D)** and lean mass **(E)** quantified by EchoMRI at the end of the diet period. **(F)** Liver weights at sacrifice after 6 hours fasting period. Statistical analysis was performed by one-way ANOVA with Tukey’s multiple comparison test (A-D) or Kruskal-Wallis with Dunn’s test (E-F). Different letters indicate significant differences (p<0.05) between groups, with common letters indicating no significant difference. Values are represented as mean ±SD.

16-week-old male B6D2F1 mice were fed the above diets for 8 days. Body weight and food consumption were tracked across the diet period (**Fig. S1, C-D**). All experimental diets resulted in significant weight loss compared to mice fed the CTL diet, with KD0P resulting in the greatest magnitude of weight loss (18.8% reduction) (**Fig. 1B**). A similar degree of weight loss was observed following PF and KD10P diets (11.7% and 10.2% reduction, respectively). All diets tended to have a lower calorie intake versus control, but only KD0P had a statistically significant decrease (**Fig. 1C**). Weight loss was primarily attributed to significant reductions in fat mass, which was observed in all diet groups, except KD20P (**Fig. 1D**). However, significant reductions in lean mass were also observed in PF and KD0P groups (**Fig. 1E**). Liver weight was also reduced following all experimental diets, with KD0P and KD5P groups significantly different from CTL mice (**Fig. 1F**).

### 3.2 Protein-free, but not ketogenic, diet improves insulin sensitivity following short-term feeding

To evaluate the metabolic effects of short-term protein or carbohydrate restriction, markers of systemic glucose homeostasis were assessed. Mice fed PF and KD diets, except for KD20P, had reduced fasting blood glucose and serum insulin after one week (**Fig. 2, A-B**). KD diets resulted in elevated fasting ketone levels, which was blunted by the addition of protein beyond 10% kilocalories (KD20P) (**Fig. 2C**). Increased ketone levels were not observed in mice fed PF.

**Figure 2.**
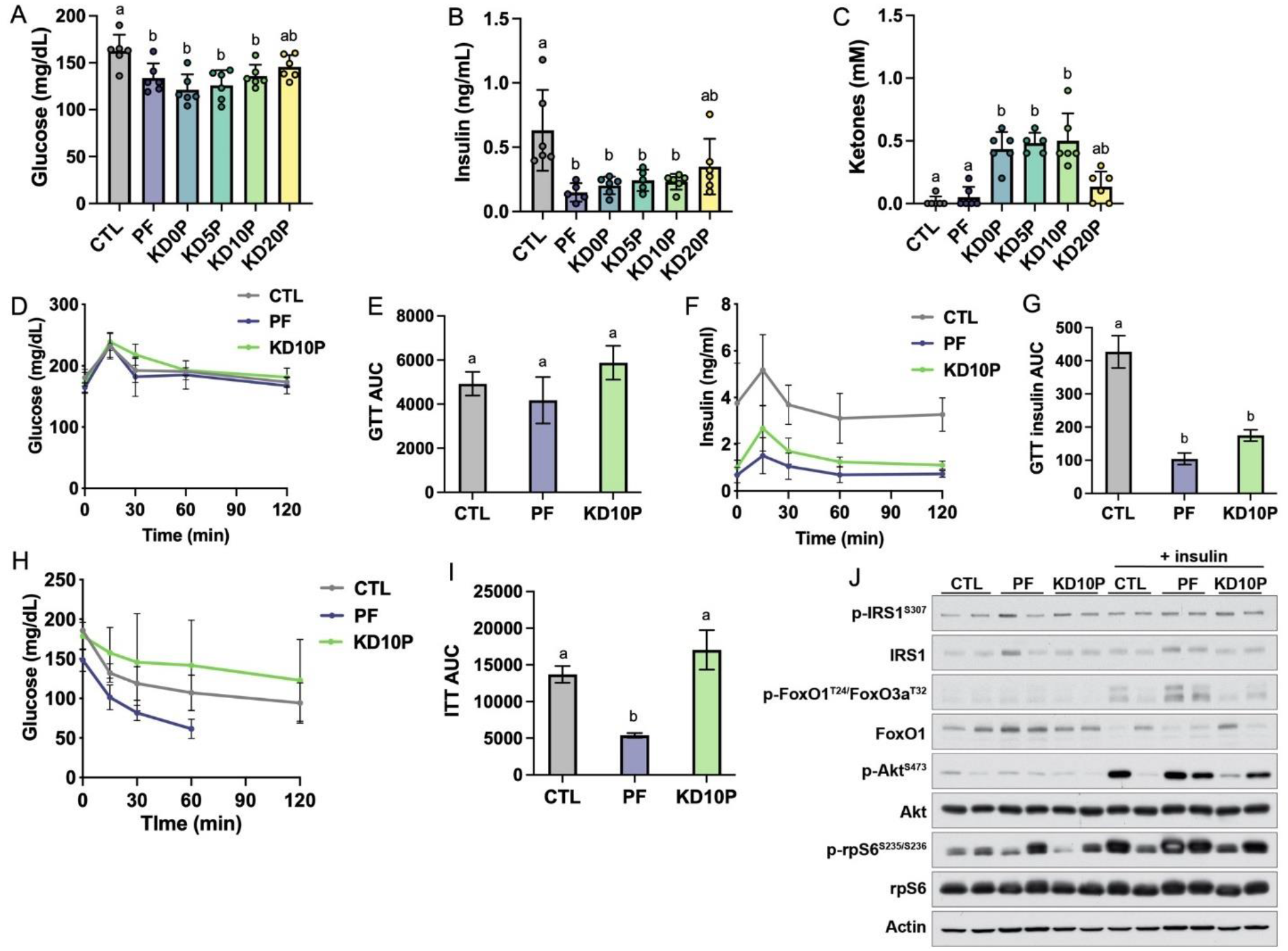
Protein-free, but not ketogenic, diet improves insulin sensitivity following short-term feeding. After 8 days of diet feeding, mice were fasted for 6 hours prior to sacrifice (n=6 per diet). Blood was collected and blood glucose **(A)** and ketones **(B)** were measured with handheld meters. **(C)** Serum insulin was measured by ELISA. **(D)** Blood glucose levels during an oral glucose tolerance test (OGTT) in a separate cohort of 6-hour fasted mice fed experimental diets for 5 days (n=4 per diet) and **(E)** corresponding area under the curve (AUC). **(F)** Serum insulin measurements collected during the OGTT and **(G)** corresponding AUC. **(H)** Blood glucose levels during an insulin tolerance test in 4-hour fasted mice fed experimental diets for 7 days (n=4 per diet) and **(I)** corresponding AUC. Measurements were halted after 60 minutes in PF-fed mice due to severe hypoglycemia. **(J)** Mice fed experimental diets for 10 days were fasted for 6 hours and then injected intraperitoneally with vehicle or insulin and sacrificed after 20 minutes. Immunoblots show insulin/Akt/mTORC1 signaling in liver lysates (n=2 per diet per treatment). Statistical analysis was performed by one-way ANOVA with Tukey’s multiple comparison test (A, E, F, I) or Kruskal-Wallis with Dunn’s test (B-C). Different letters indicate significant differences (p < 0.05) between groups, with common letters indicating no significant difference. Values are represented as mean ±SD.

Glucose and insulin tolerance were further assessed in a separate cohort of mice fed CTL, PF, or KD10P diets (n=4 per diet). Neither PF nor KD10P diets significantly affected glucose tolerance (**Fig. 2D-E; Fig. S2A-B**). Despite the lack of significant change in glucose tolerance, serum insulin concentrations during the GTT were significantly lower in both PF-and KD-fed mice (**Fig. 2F-G**), suggesting that PF and KD10P may increase insulin sensitivity. However, only PF-fed mice demonstrated improved insulin tolerance, while KD10P-fed mice tended to have worse insulin tolerance, compared to CTL-fed mice (**Fig. 2H-I; Fig. S2C-D**). To evaluate whether PF or KD10P diets enhanced the hepatic response to insulin, insulin/Akt signalling was assessed in liver tissue following intraperitoneal insulin injection. Phosphorylation of Akt and its target, FoxO1, tended to increase in response to insulin in livers from PF compared to CTL mice (**Fig. 2J; Fig. S2E-F**). Insulin/Akt signalling tended not to be enhanced in livers from KD10P compared to CTL mice.

### 3.3 Protein-free, but not ketogenic, diets induce molecular markers of dietary protein restriction

Next, we aimed to determine whether ketogenic diets induce established markers of protein restriction, including serum FGF21 levels, hepatic *Fgf21* and *Asns* mRNA expression and brown adipose tissue (BAT) *Ucp1* mRNA expression. Serum FGF21 was increased ~5-fold in mice fed either protein-free diets, PF or KD0P, compared to CTL mice (**Fig. 3A**). Serum FGF21 was also significantly increased in mice fed KD5P, but to a lesser extent, and was not increased in mice fed either KD10P or KD20P, compared to CTL mice.

**Figure 3.**
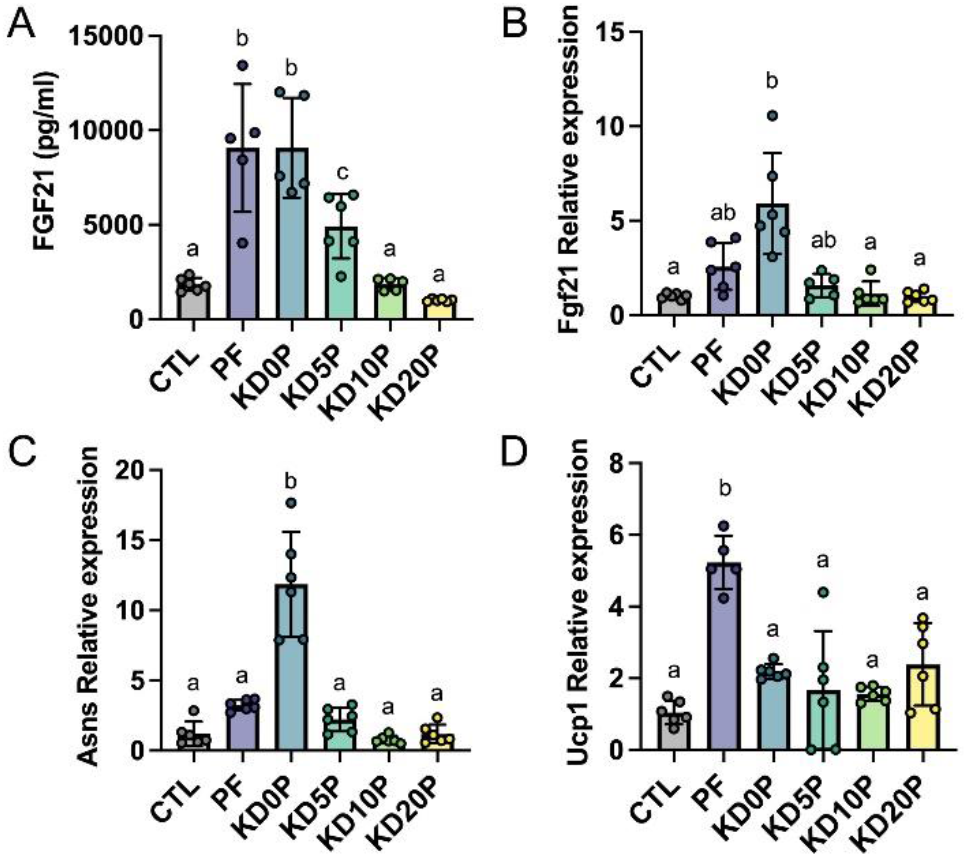
Protein-free, but not ketogenic, diets induce molecular markers of dietary protein restriction. **(A)** Serum FGF21 was measured in 6-hour fasted mice fed experimental diets for 8 days. Liver gene expression of *Fgf21* **(B)** and *Asns* **(C)** and brown adipose tissue expression of *Ucp1* **(D)** were measured by RT-qPCR and normalized to actin expression. Statistical analysis was performed by one-way ANOVA with Tukey’s multiple comparison test (A, C-D) or Kruskal-Wallis with Dunn’s test (B). Different letters indicate significant differences (p < 0.05) between groups, with common letters indicating no significant difference. Values are represented as mean ±SD.

The adaptation to dietary protein restriction includes upregulation of activating transcription factor 4 (ATF4) gene targets *Fgf21* and *Asns* in the liver, as well as upregulation of *Ucp1* expression in BAT. Next, we compared the induction of these gene signatures by PF and KD diets. As expected, liver *Fgf21* and *Asns* expression was elevated in mice fed PF (**Fig. 3B-C**). However, neither *Fgf21* nor *Asns* expression was increased in the liver of mice fed KDs containing at least 5% protein (**Fig. 3B-C**). In agreement with previous reports, BAT *Ucp1* expression was significantly elevated in mice fed PF compared to those fed CTL (~5-fold, **Fig. 3D**). While BAT *Ucp1* expression was slightly increased in mice fed either of the KD diets compared to those fed the CTL diet, none of these differences were significant. Interestingly, although mice fed the protein-free KD, KD0P, demonstrated significantly greater elevations in hepatic *Fgf21* (~2-fold difference) and *Asns* expression (~4-fold difference) compared to mice fed the carbohydrate-rich PF diet, they showed only equivalent or less circulating FGF21 and BAT *Ucp1* expression (**Fig. 3A-D**).

### 3.4 Protein-free and ketogenic diets produce distinct hepatic transcriptomic signatures

To further explore changes in hepatic gene expression as a function diet, we performed RNA sequencing. Data reduction by multidimensional scaling showed distinct clustering of samples by group with KD20P showing the least divergence from CTL and KD0P showing the greatest divergence (**Fig. 4A**). Key genes in the amino acid restriction response showed a greater relative increase in KD0P vs PF, and no response in KD10P, including *Fgf21, Asns* and *Psat1* (**Fig. 4B-D**). Interestingly, although *Atf4* expression was similarly elevated in KD0P compared to PF livers, it also tended to increase across the other KD groups (**FigS3A**), suggesting it may not be a key driver of this response. When weighted gene correlation network analysis (WGCNA) was performed, a single module containing these three genes (*Fgf21, Psat1, Asns*) was identified. This module contained 51 genes which exhibited similar behavior across groups including a proportionally greater increase in KD0P vs PF (**Fig. S3B**). Several other genes characteristic of the amino acid stress response were identified in this module including tRNA synthetases (*Yars, Cars, Mars, Farsa, Rars*), *Pck2* and *Mthfd2*. When transcription factor (TF) binding site analysis was performed, binding motifs for Ventx, Thap11, Atf4 and Sox21 were the top enriched hits compared to a background list of 1000 random genes (**Fig. S4C**). Binding motifs for multiple FOX family TFs were significantly depleted in this module, including Foxo4, Foxo6 and Foxl1 (**Fig. S3C**).

**Figure 4.**
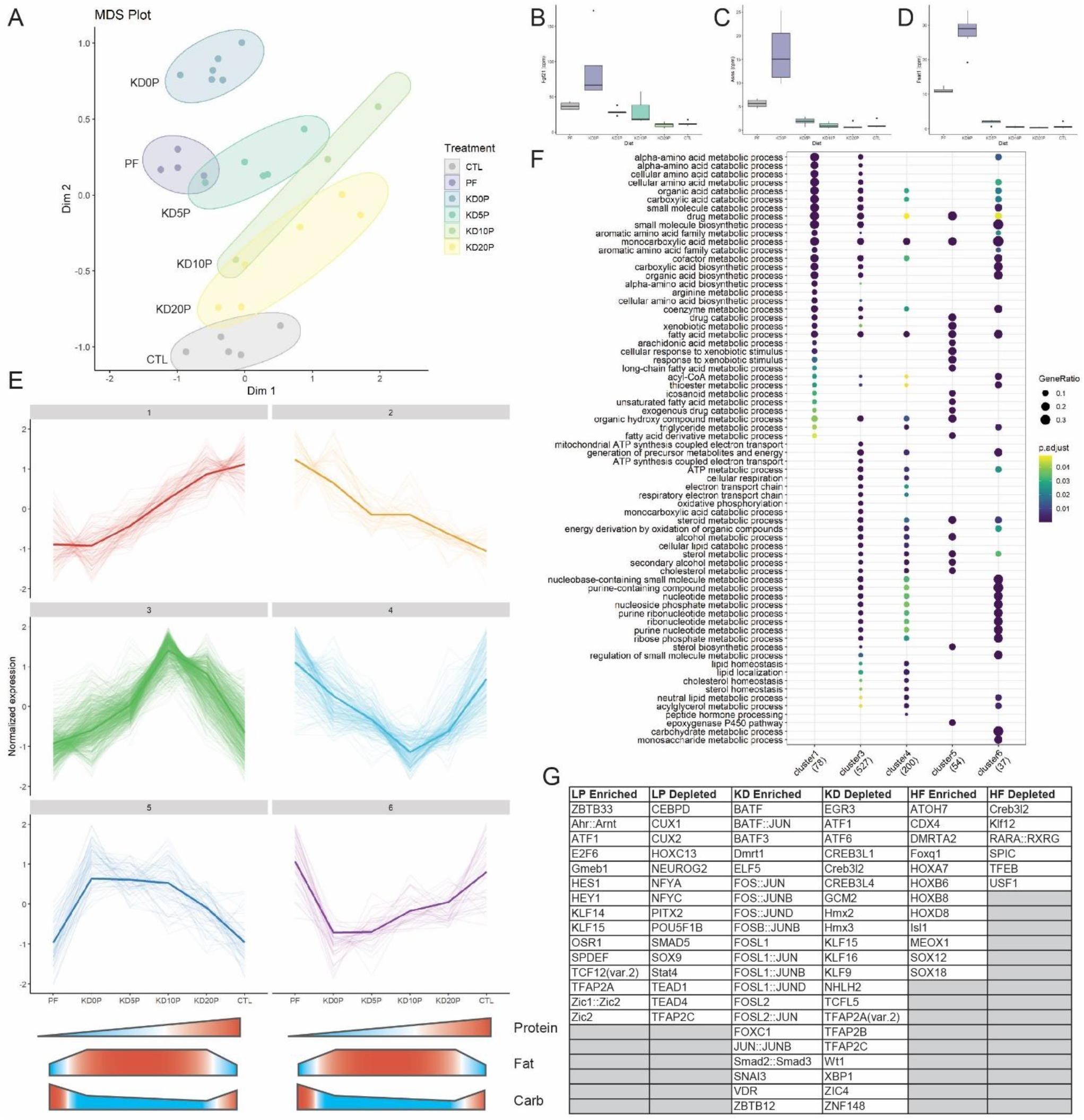
Protein-free and ketogenic diets produce distinct hepatic transcriptomic signatures. **(A)** Multidimensional scaling (MDS) plot across all diet groups/samples. **(B)** Expression of amino acid responsive genes Fgf21, **(C)** Asns and **(D)** Psat1 presented at counts per million (cpm). **(E)** Cluster analysis of the top 10% of most variable genes in the dataset. Clusters 1 and 2 represent protein-responsive genes. Clusters 3 and 4 represent genes responding to classic ketogenic diet. Clusters 5 and 6 include genes responsive to dietary fat content. **(F)** Pathway over-representation analysis of genes from the clusters identified in E. **(G)** Enriched and depleted transcription factor (TF) binding motifs in the clusters identified in E. LP enriched includes TF binding motifs which are significantly depleted in cluster 1 and significantly enriched in cluster 2. The same method was used for KD in clusters 3 and 4 and high fat (HF) in clusters 5 and 6.

To better understand how gene networks change in response to the experimental diets, unsupervised clustering of the most variable 10% of genes in the dataset was performed. Six major clusters were identified from this analysis (**Fig. 4E**). Clusters 1 and 2 responded to dietary protein content by linearly decreasing or increasing respectively. Clusters 3 and 4 were either increased or decreased respectively by the classic ketogenic diet (KD10P). Clusters 5 and 6 were comprised of genes which increased or decreased respectively as a function of increased dietary fat content, largely independent of ketosis.

Gene set overrepresentation analysis showed that the genes linearly responding to protein content were largely involved in amino acid metabolic processes, fatty acid metabolism and CoA metabolic processes (**Fig. 4F**). Cluster 3 (increasing with classic KD) was enriched for genes involved in oxidative phosphorylation, lipid catabolism and nucleotide metabolic processes, which were not observed in the protein responsive clusters. There was some overlap between clusters 1 and 3 regarding amino acid catabolic processes. The pathways increased in response to increased dietary fat content (cluster 5) were primarily involved in lipid metabolism and CYP regulation. Interestingly, pathways enriched in genes that decreased in response to dietary fat tended to overlap with those that were increased in response to classic KD, particularly those involving nucleotide metabolism.

To investigate TFs that may be regulating these differential responses to diet, binding site motif analysis was performed across the clusters. To determine TFs that may be enriched in response to low protein, the overlap of TFs significantly depleted in cluster 1 and significantly enriched in cluster 2 was taken. A similar approach was taken to identify TFs enriched by KD using clusters 3 and 4, and TFs enriched by high fat using clusters 5 and 6. In the low protein condition, motifs for Atf1, Ahr::Arnt and E2F6 binding were among the most enriched (**Fig. 4G, Fig. S3D-E**). In the KD condition, binding sites for FOS and JUN family proteins were highly enriched including Fos, Fosb, Fosl1 and Fosl2 (**Fig. 4G, Fig. S3F-G**). In the high fat condition, binding motifs for HOX and SOX family proteins were significantly enriched, independent of ketogenesis (**Fig. 4G, Fig. S4H-I**).

## 4 Discussion

This study utilized six distinct diets to dissect the influence of protein content on the short-term metabolic adaptations to the high-fat, low-carbohydrate ketogenic diet. Mice fed a protein-free diet (PF) or one of three ketogenic diets with 10% or less protein lost weight, which was primarily attributed to loss of fat mass, and had reduced fasting blood glucose and insulin levels. Despite similar physiological effects of KD and PF diets, the diets differed with respect to some metabolic and molecular markers. While mice fed PF demonstrated improved insulin sensitivity and liver insulin signaling, mice fed KD did not. With the exception of the protein-free KD (KD0P), KD diets did not lead to elevations in serum or tissue markers of protein restriction, including circulating FGF21, hepatic *Fgf21* and *Asns* and BAT *Ucp1* expression. Furthermore, PF and classic KD diets elicited distinct changes in global gene expression. These data indicate that KD does not induce a molecular response to protein restriction and that PF and KD diets impact health and metabolism via different underlying mechanisms.

Defining the effects of KD on metabolic health has been complicated by discrepancies in diet composition across studies, particularly in studies which use “control chow” diets. KD diets in published rodent studies contain between 4.5-10% protein with feeding durations from 3 days to 14 months. The present study helps clarify discrepancies by using standardized semi-purified diets that control for micronutrient composition and span the range of protein content found in published KD diets. In addition to KD diet composition, published studies differ with respect to control diet formulation. For studies that utilize a semi-purified ketogenic diet, the control diet should also be semi-purified in order to ensure that differences in micronutrient content or nutrient source do not account for any of the phenotypic observations attributed to macronutrients. The diets used in the present study were prepared from the same semi-purified ingredients in order to evaluate the precise contribution of individual macronutrients.

It is also important to note that some effects of KD may be specific to the duration of feeding. Our finding that mice fed KD for 8 days are insulin resistant is aligned with other studies of short-term KD (12, 13, 55). However, studies in which mice were fed KD for 1 month (11) or 60 weeks (6) found improved insulin sensitivity, indicating additional adaptations to longer-term feeding. This study aimed to assess the physiological and molecular response to short-term feeding, which will be important for defining the potential therapeutic benefits of KD for recent cancer and weight loss indications.

The present study found no elevations in serum FGF21 and no induction of hepatic *Fgf21* expression in mice fed the classic KD containing 10% protein. While some published studies report increased plasma or hepatic FGF21 after 4-5 weeks (12, 55), the KD used in these studies contained approximately 5% protein. Thus, our finding that FGF21 was only induced in response to KD containing 5% or less protein is consistent with these reports and demonstrates that the effects of ketogenesis can be disconnected from those of FGF21. Furthermore, by carefully titrating the levels of protein in KD, we demonstrate that circulating FGF21 is induced in response to decreasing dietary protein content, further establishing it as a marker of protein restriction. This result is consistent with a published report demonstrating that low protein, high carbohydrate diets are associated with maximal FGF21 induction (56) and demonstrates that KD may act independently of dietary protein restriction to elicit metabolic or other health benefits. Therefore, we propose that future studies on KD utilize diets containing at least 10% protein in order to ensure that observations are due to carbohydrate restriction, rather than protein restriction.

FGF21 signaling in the central nervous system (CNS) has contributes to increased food intake during low-protein diet (5% protein) feeding, a phenomenon known as protein leverage (57). However, FGF21 signaling is not sufficient to overcome reduced food intake and body weight in response to very-low-protein diets (1% protein). This decrease in intake is at least partially attributed to reduced hypothalamic mTOR signaling (58, 59). Interestingly, increased BAT *Ucp1* expression requires intact FGF21 signaling in the brain, but not in the adipose tissue (57). In our study, mice fed the high-fat, protein-free diet (KD0P) had equal serum FGF21 and hepatic *Fgf21* expression levels vs high-carbohydrate PF-fed mice. However, the KD0P group had significantly reduced energy intake and did not have elevated BAT *Ucp1* expression compared to mice fed CTL and PF diets. Given that brain FGF21 signaling promotes energy intake and is required for *Ucp1* induction, it is possible that these characteristics of mice fed the high-fat KD0P are a result of FGF21 resistance, which has been associated with high-fat diet-induced obesity in mice (60). This may also explain why markers of protein restriction are induced in response to KD0P in the liver, but not BAT.

Consistent with this observed difference, on the transcriptional level PF and KD drove divergent gene expression programs. First, the overall transcriptional response to PF diet was amplified by substituting carbohydrate for fat (PF vs KD0P, **Fig. 4A**). The high fat background drove higher expression of multiple genes involved in amino acid synthesis including *Psat1, Phgdh* and *Asns*, as well as the amino acid responsive transcription factor *Atf4*. WGCNA analysis identified a module containing these amino acid responsive genes which were further increased by high fat background. Atf4 was the third-most enriched TF-binding motif among these genes. However, *Atf4* transcript level also increased in KDs where these amino acid-related targets did not increase. This leaves an open question about the role of ATF4 in driving this differential expression in response to KD0P vs PF.

Clustering analysis reveal expression modules which were responsive to protein, to classic KD (KD10P) and to fat content independent of protein or ketogenesis. Interestingly, the largest number of genes were positive responders to classic KD, with over 500 genes clustering in this pattern. The protein responsive clusters were primarily enriched for genes involved in amino acid metabolism and fatty acid metabolism, with increased expression of genes involved in non-essential amino acid synthesis and lipid degradation pathways. Among the most enriched pathways in the classic KD responsive gene set was oxidative phosphorylation, with expression of many mitochondrial genes, such as *Ndufa10, Sdhb* and *Sdhc* increased in response to KD10P. Interestingly, these genes tended to be suppressed by reduced protein, even in the high fat background. Similarly, nucleotide metabolic processes were also enriched in response to classic KD (cluster 3). These pathways show some overlap with mitochondrial-related pathways and include genes like *Mpc2*, *Upp2* and *Pmvk*. Many of these nucleotide-related processes tended to show an overall suppression by high fat diet, suggesting that these increases are specific to the classic ketogenic diet, and not just due to high fat content. TF analyses showed enrichment of Atf and Hif signaling in low protein groups, with Atf1, Atf7 and Hif/Arnt showing strong enrichments. In the KD clusters, binding motifs for Fos/Jun were robustly enriched. These differential TF signatures support the divergent expression profiles seen in the gene expression analysis.

While this study clarifies that protein restriction shows relatively little overlap with KD, multiple outstanding questions remain. In particular, it will be important to understand how the molecular and metabolic adaptations to KD differ across feeding duration and how short versus long-term adaptations are connected to stress resistance, weight loss and glucose homeostasis. In addition, it is unknown how ketosis or other KD-induced alterations impart the associated physiological changes. The transcriptomics data presented here provide novel insight into potential underlying mechanisms of KD and can serve as a starting point for future mechanistic studies. In conclusion, KD and protein-restricted diets produce similar physiological outcomes, though through distinct mechanisms, and more work is needed to define how short-term KD may be utilized therapeutically.

## 6 Author Contributions

J.M., M.M., S.M., and K.K. conceived of and designed the study. K.K., M.M., and S.M. performed experiments. K.K. and M.M. analyzed and interpreted the data. M.M. performed transcriptomic analyses. K.K. and M.M. drafted the manuscript, which was reviewed by S.M.

## 8 Acknowledgments

We thank members of the Mitchell Lab for critical discussions on study design and data. We also thank Brendan Manning for support and input during manuscript drafting.

## 9 Data Availability Statement

The RNA seq datasets generated for this study can be found on the NIH Sequence Read Archive (SRA) under the BioProject PRJNA777819: Profiling the hepatic transcriptional response to low protein versus low carbohydrate diets. A companion web-based application for data exploration is published at macarthur.shinyapps.io/kdshiny.

## 11 Supplementary Figures

**Supplementary Figure 1.**
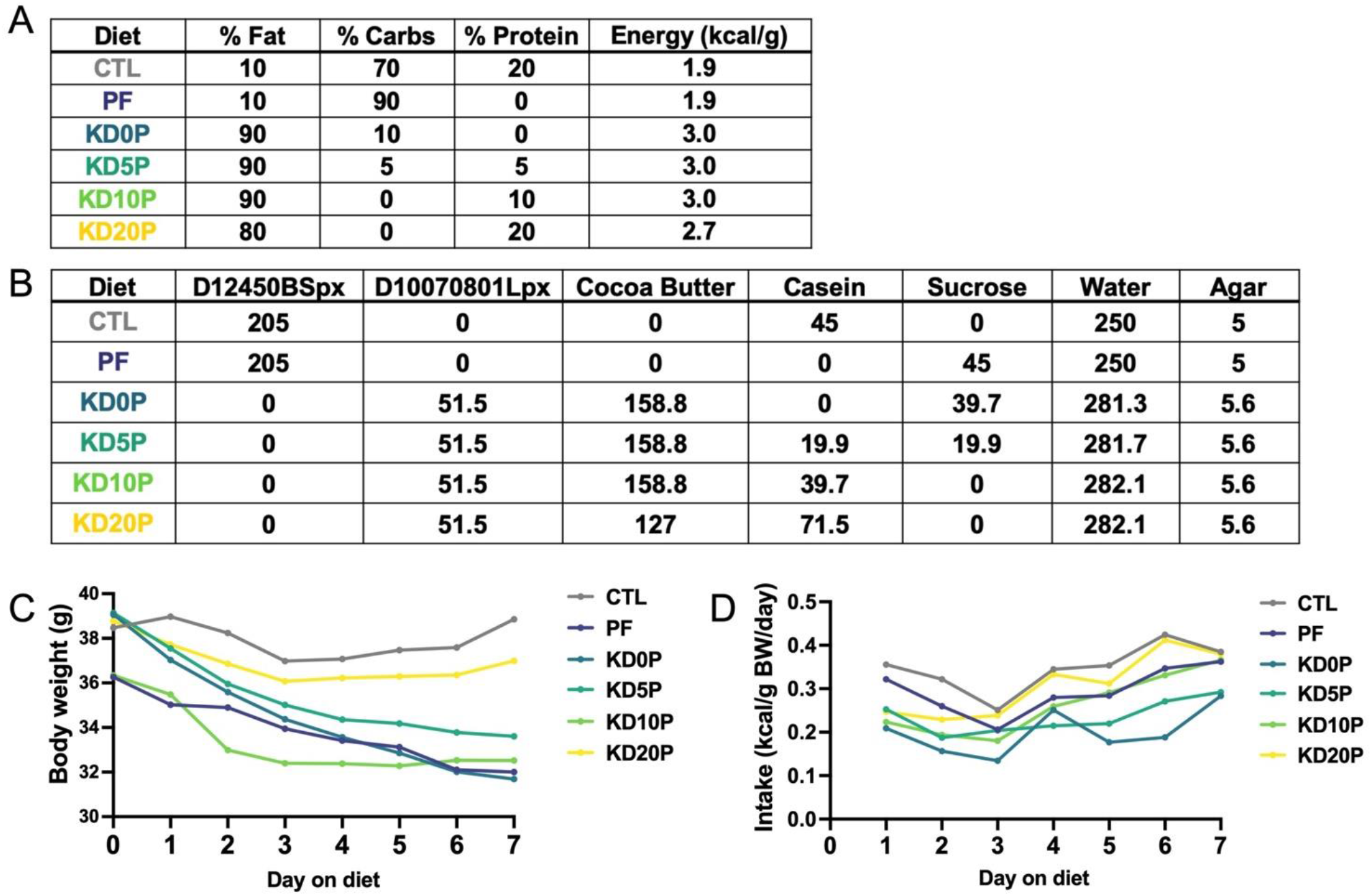
**(A)** Table of macronutrient compositions and energy densities of the experimental diets. **(B)** Table of ingredients in experimental diets in grams. **(C)** Body weights in grams across 7 days of feeding experiments diets. **(D)** Energy intakes in kcals per gram body weight per day across 7 days of feeding experimental diets. Intake was normalized to the summed body weight of 2 mice co-housed per cage and presented per cage.

**Supplementary Figure 2.**
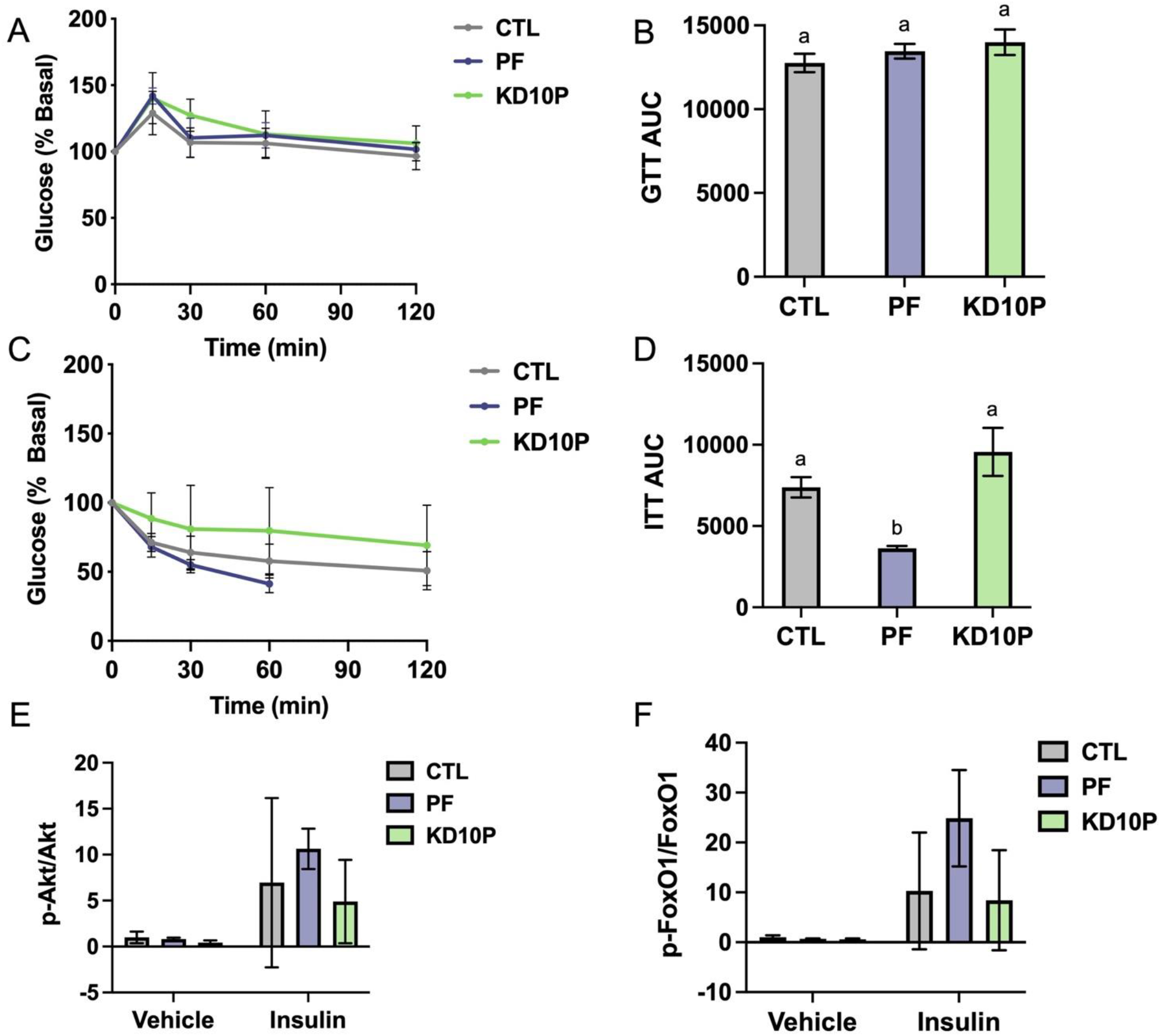
**(A)** Blood glucose levels during oral glucose tolerance test, normalized to baseline and **(B)** corresponding area under the curve (AUC). **(C)** Blood glucose levels during insulin tolerance test normalized to baseline and **(D)** corresponding AUC. **(E)** Quantification of phospho-AKT normalized to total AKT by western blot. **(F)** Quantification of phospho-FoxO1 normalized to total FoxO1 by western blot.

**Supplementary Figure 3.**
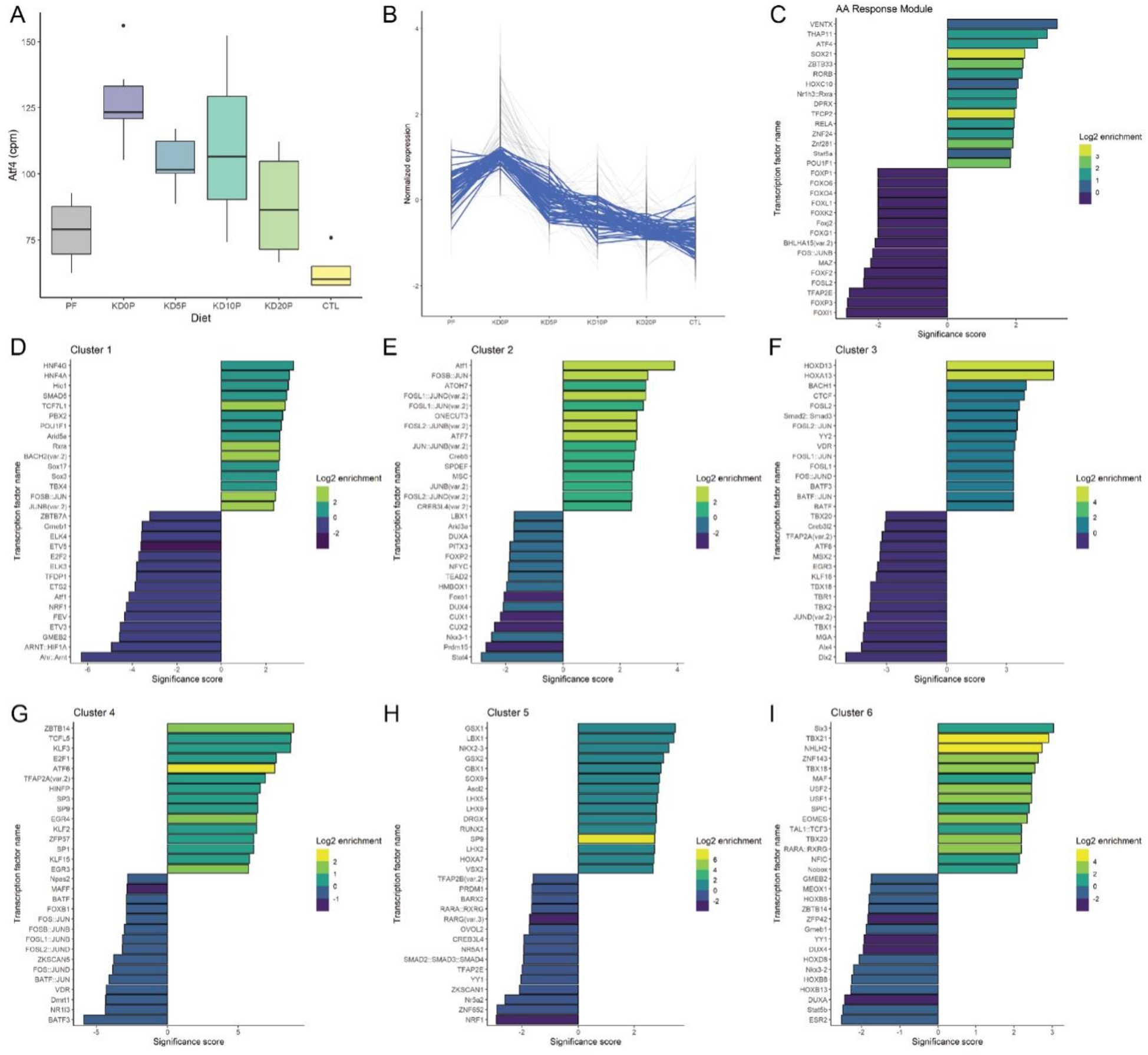
**(A)** Hepatic Atf4 transcript levels as counts per million (cpm) across experimental diets. **(B)** Weighted gene correlation network analysis module containing amino acid responsive genes including Psat1, Asns and Fgf21. **(C)** Top 10 enriched and depleted transcription factor binding motifs corresponding to the module in B. **(D-I)** Top 10 enriched and depleted transcription factor binding motifs in clusters 1 through 6.

**Table S1.** Composition of low fat base diet.

| Product # | D12450BS |  | D12450BSpx |  |
| --- | --- | --- | --- | --- |
| % | gm | kcal | gm | kcal |
| Protein | 19.2 | 20.0 | 0.0 | 0.0 |
| Carbohydrate | 67.3 | 70.0 | 83.3 | 87.5 |
| Fat | 4.3 | 10.0 | 5.3 | 12.5 |
| Total |  | 100.0 |  | 100.0 |
| kcal/gm | 3.85 |  | 3.81 |  |
| Ingredient | gm | kcal | gm | kcal |
| Casein, 80 Mesh | 0 | 0 | 0 | 0 |
| Casein, Enzyme Hydrolyzed | 200 | 800 | 0 | 0 |
| L-Cystine | 3 | 12 | 0 | 0 |
| Sucrose | 350 | 1400 | 350 | 1400 |
| Maltodextrin 42 | 350 | 1400 | 350 | 1400 |
| Cellulose, BW200 | 40 | 0 | 40 | 0 |
| Xanthan Gum | 10 | 0 | 10 | 0 |
| Soybean Oil | 25 | 225 | 25 | 225 |
| Lard | 20 | 180 | 20 | 180 |
| Mineral Mix S10011 | 0 | 0 | 0 | 0 |
| Mineral Mix S10026 | 10 | 0 | 10 | 0 |
| DiCalcium Phosphate | 13 | 0 | 13 | 0 |
| Calcium Carbonate | 5.5 | 0 | 5.5 | 0 |
| Potassium Citrate, 1 H<sub>2</sub>O | 16.5 | 0 | 16.5 | 0 |
| Vitamin Mix V10001 | 10 | 40 | 10 | 40 |
| Choline Bitartrate | 2 | 0 | 2 | 0 |
| FD&C Yellow Dye #5 | 0.05 | 0 | 0.05 | 0 |
| **Total** | **1055.05** | **4057** | **852.05** | **3245** |

**Table S2.** Composition of high fat base diet.

Water Suspendible Rodent Diet With 90 kcal% Fat (Mostly Cocoa Butter)
| Product # | D10070801 |  | D10070801L |  | D10070801Lpx |  |
| --- | --- | --- | --- | --- | --- | --- |
|  | gm% | kcal% | gm% | kcal% | gm% | kcal% |
| Protein | 17 | 10 | 17 | 10 | 0 | 0 |
| Carbohydrate | 0 | 0 | 0 | 0 | 0 | 0 |
| Fat | 67 | 90 | 67 | 90 | 20 | 100 |
| Total |  | 100 |  | 100 |  | 100 |
| kcal/gm | 6.7 |  | 6.7 |  | 1.8 |  |
| **Ingredient** | **gm** | **kcal** | **gm** | **kcal** | **gm** | **kcal** |
| Casein | 100 | 400 | 100 | 400 | 0 | 0 |
| L-Cystine | 1.5 | 6 | 1.5 | 6 | 0 | 0 |
| Corn Starch | 0 | 0 | 0 | 0 | 0 | 0 |
| Maltodextrin 10 | 0 | 0 | 0 | 0 | 0 | 0 |
| Sucrose | 0 | 0 | 0 | 0 | 0 | 0 |
| Cellulose, BW200 | 50 | 0 | 40 | 0 | 40 | 0 |
| Xanthan Gum | 0 | 0 | 10 | 0 | 10 | 0 |
| Soybean Oil | 25 | 225 | 25 | 225 | 25 | 225 |
| Lard | 0 | 0 | 0 | 0 | 0 | 0 |
| Cocoa Butter | 381 | 3429 | 381 | 3429 | 0 | 0 |
| Primex | 0 | 0 | 0 | 0 | 0 | 0 |
| Mineral Mix, S10026 | 10 | 0 | 10 | 0 | 10 | 0 |
| DiCalcium Phosphate | 13 | 0 | 13 | 0 | 13 | 0 |
| Calcium Carbonate | 5.5 | 0 | 5.5 | 0 | 5.5 | 0 |
| Potassium Citrate, 1 H2O | 16.5 | 0 | 16.5 | 0 | 16.5 | 0 |
| Vitamin Mix, V10001 | 0 | 0 | 0 | 0 | 0 | 0 |
| Vitamin Mix, V10001C, 10x vitamins | 1 | 0 | 1 | 0 | 1 | 0 |
| Choline Bitartrate | 2 | 0 | 2 | 0 | 2 | 0 |
| Cholesterol | 0 | 0 | 0 | 0 | 0 | 0 |
| Sodium Cholic Acid | 0 | 0 | 0 | 0 | 0 | 0 |
| FD&C Yellow Dye #5 | 0.025 | 0 | 0.025 | 0 | 0.025 | 0 |
| FD&C Red Dye #40 | 0.025 | 0 | 0.025 | 0 | 0.025 | 0 |
| FD&C Blue Dye #1 | 0 | 0 | 0 | 0 | 0 | 0 |
| **Total** | **605.55** | **4060** | **605.55** | **4060** | **123.05** | **225** |

MacArthurM02.for

